## Supplementary figures and images for "Heterozygous expression of a *Kcnt1* gain-of-function variant has differential effects on SST- and PV-expressing cortical GABAergic neurons"

### Extended Figure 2-1

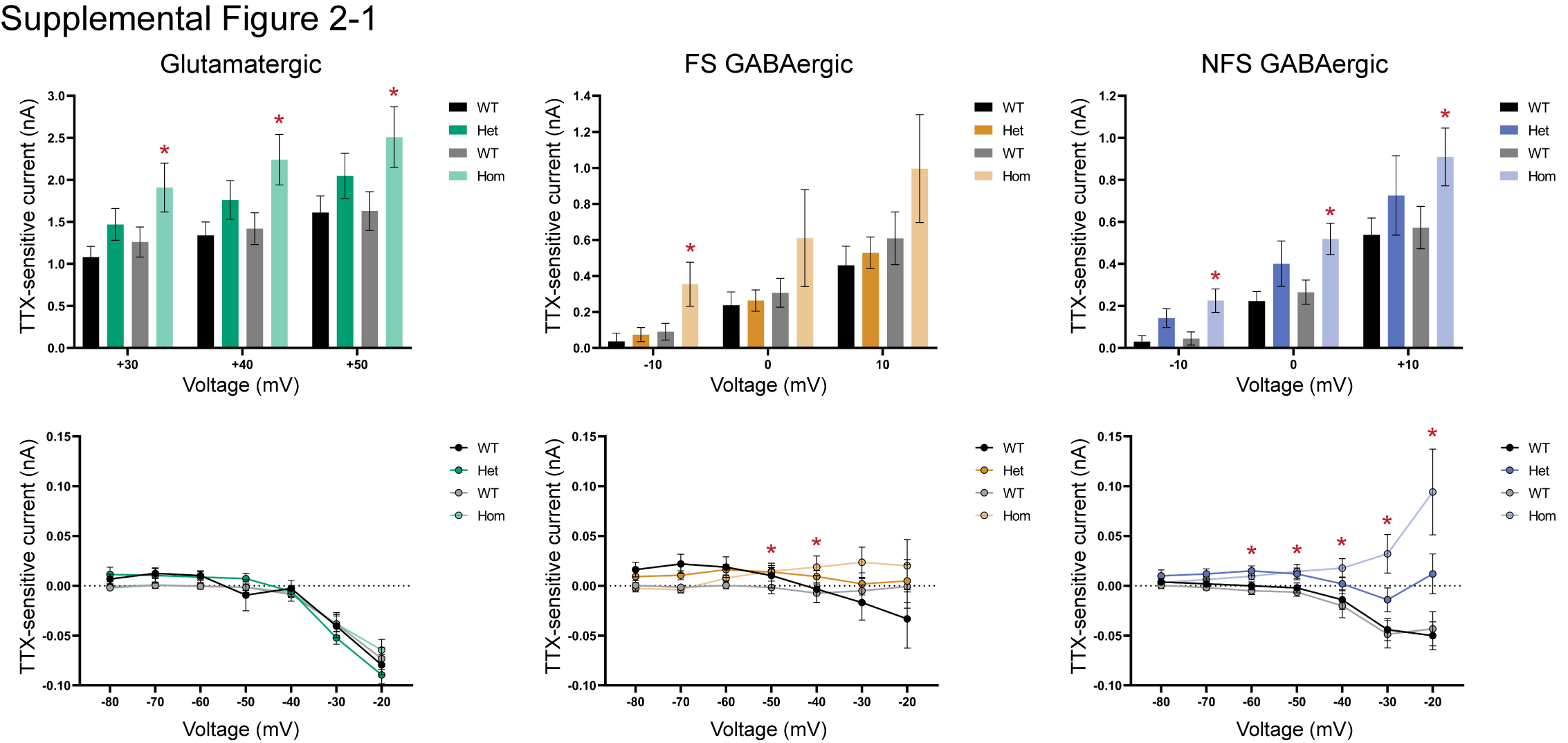

### Extended Figure 4-1

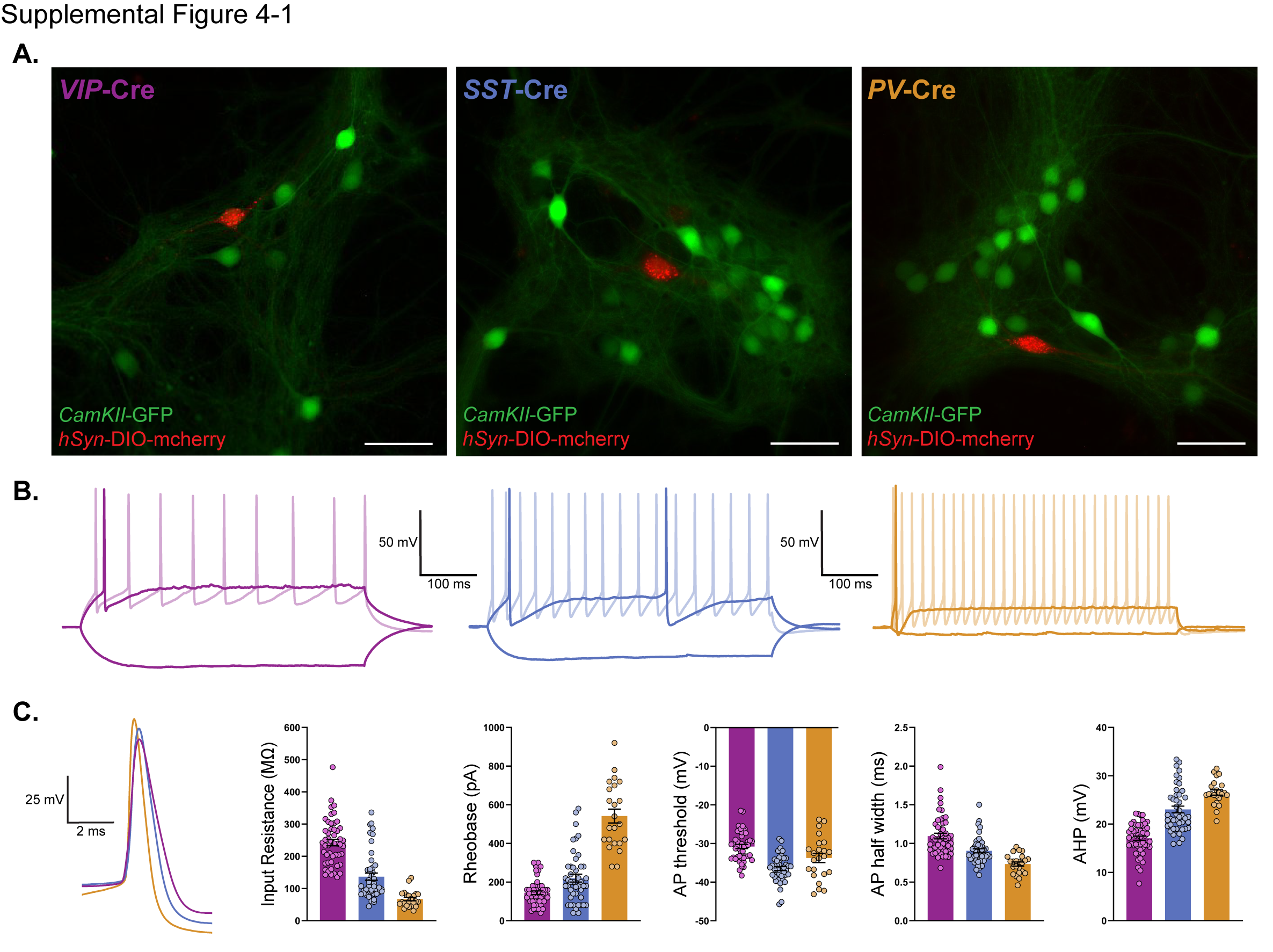

### Extended Figure 4-2

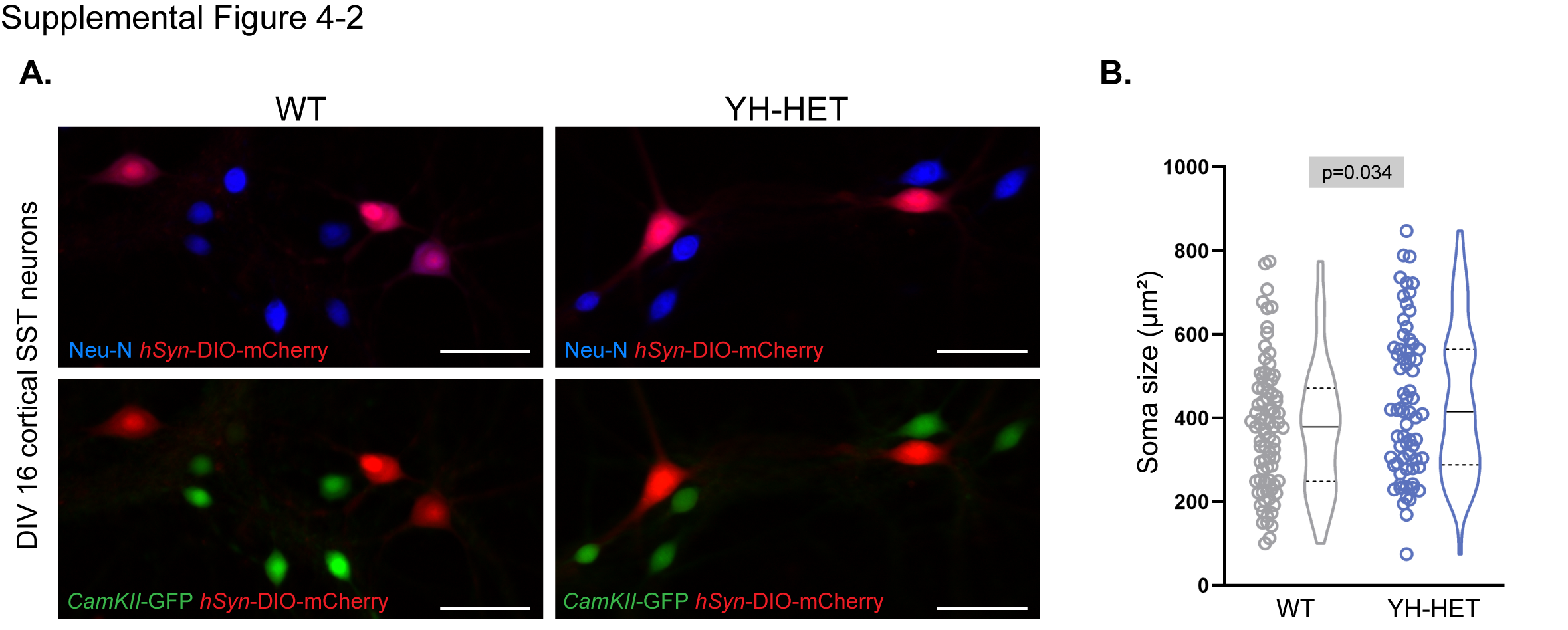

### Extended Figure 4-3

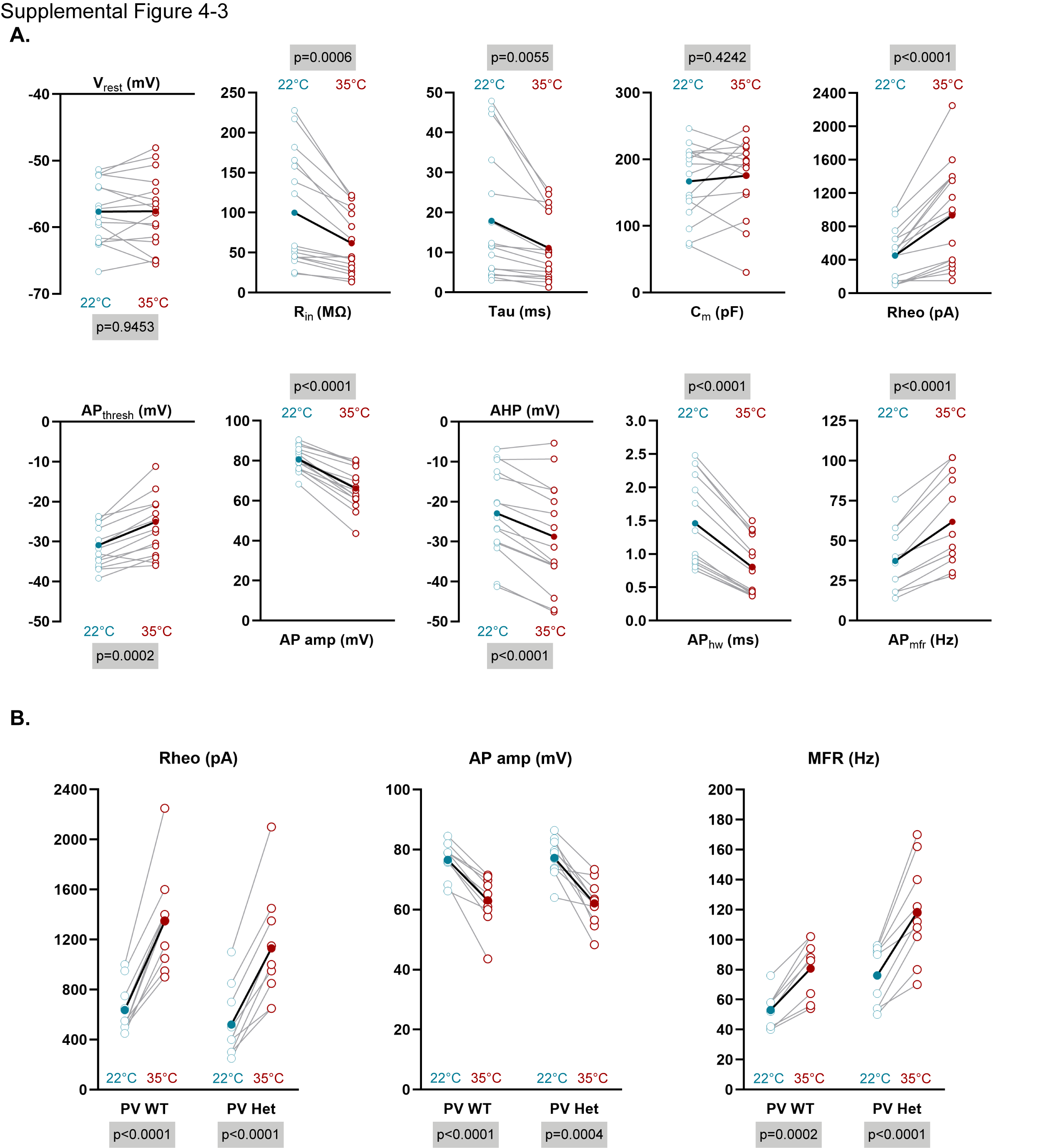

### Extended Figure 5-1

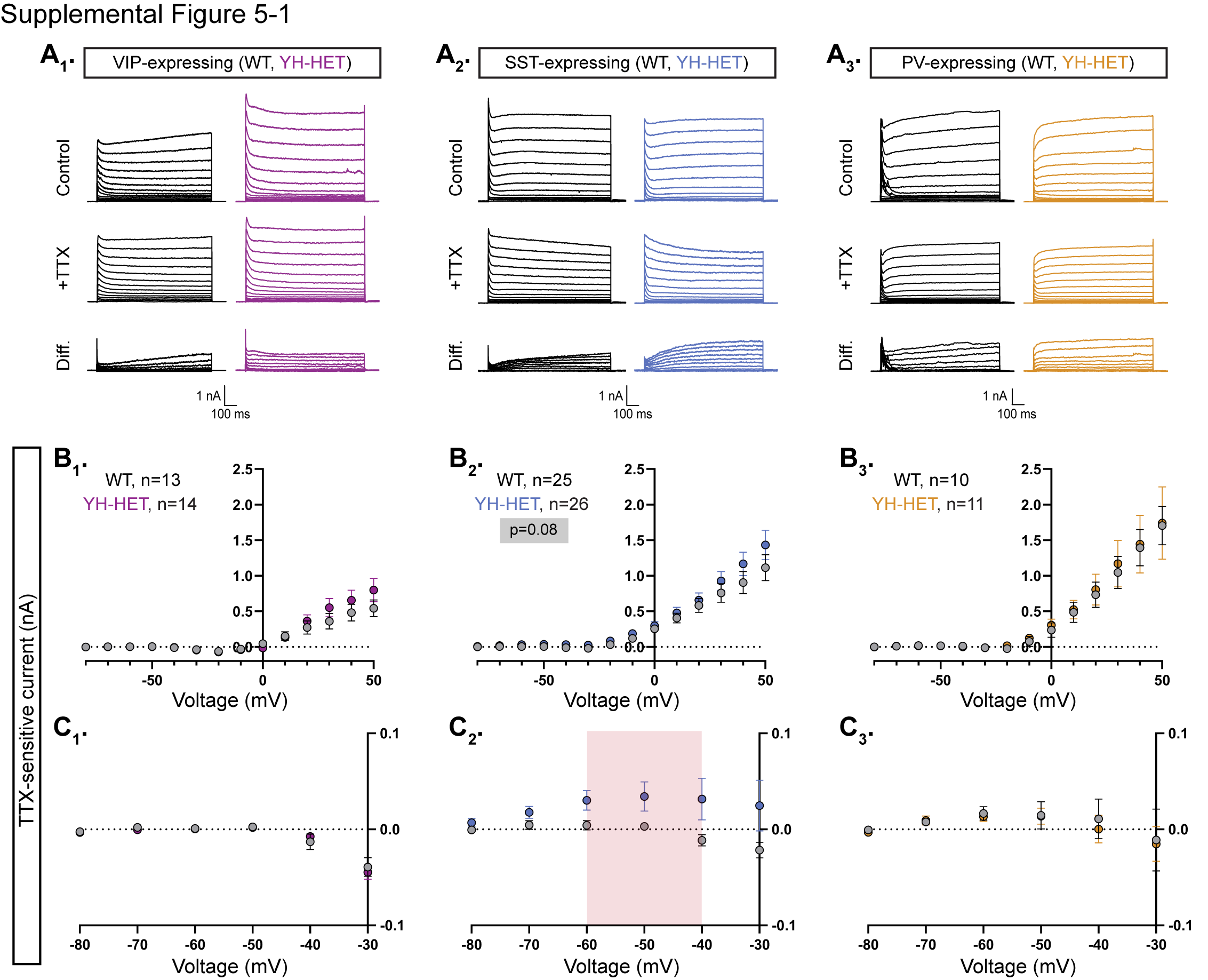

### Extended Figure 5-2

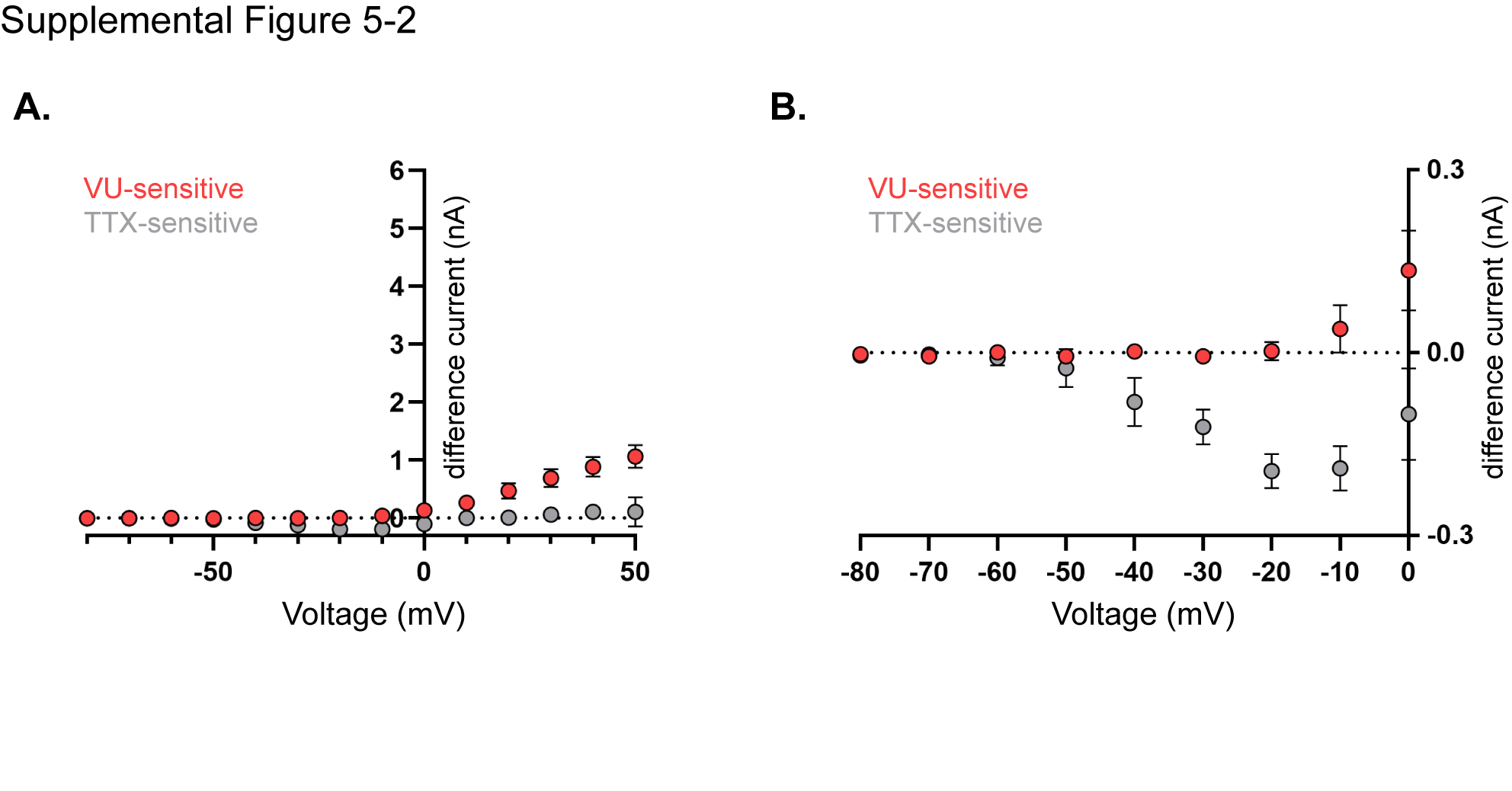

### Extended Figure 6-1

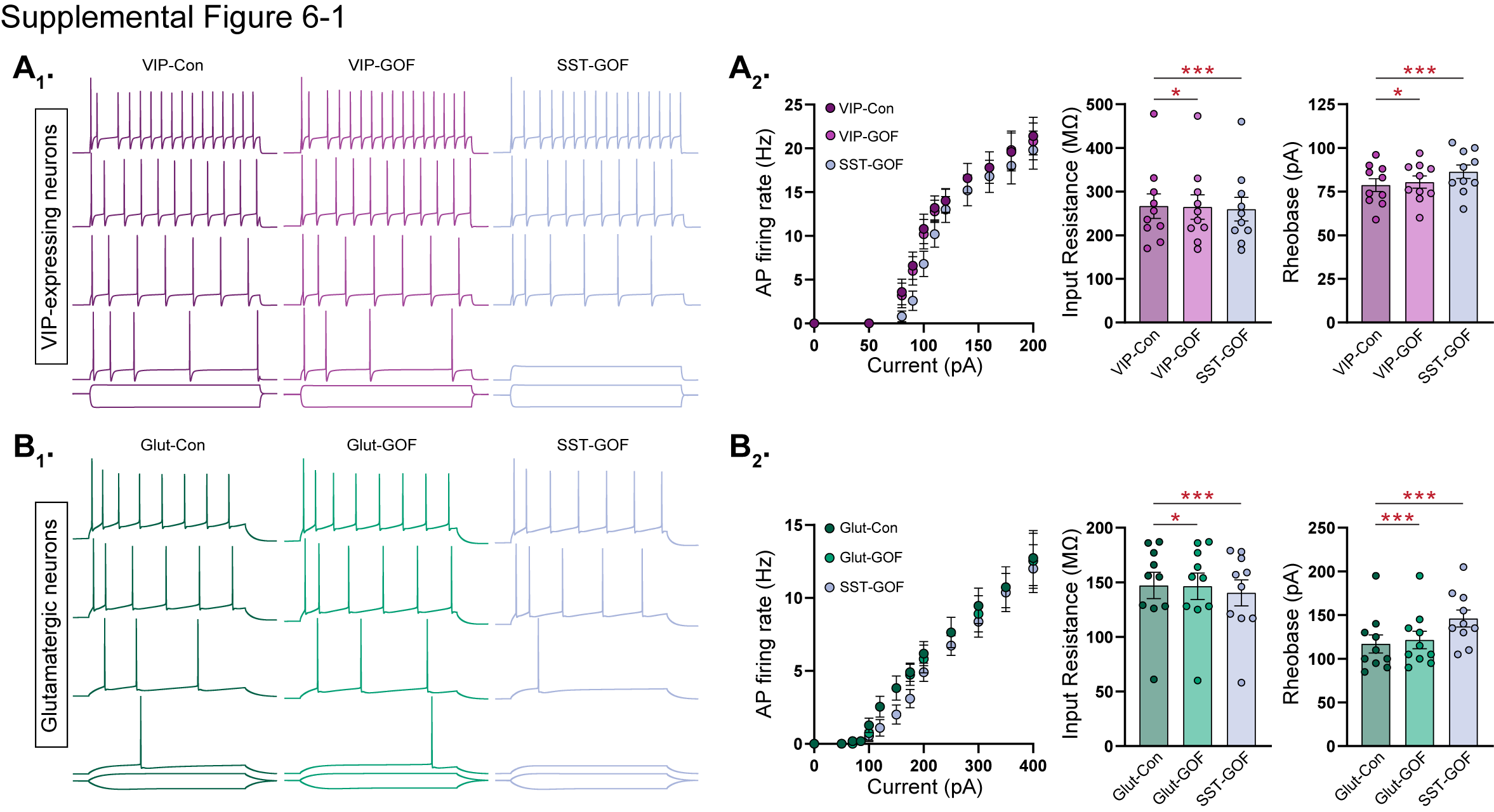

### Extended Figure 6-2

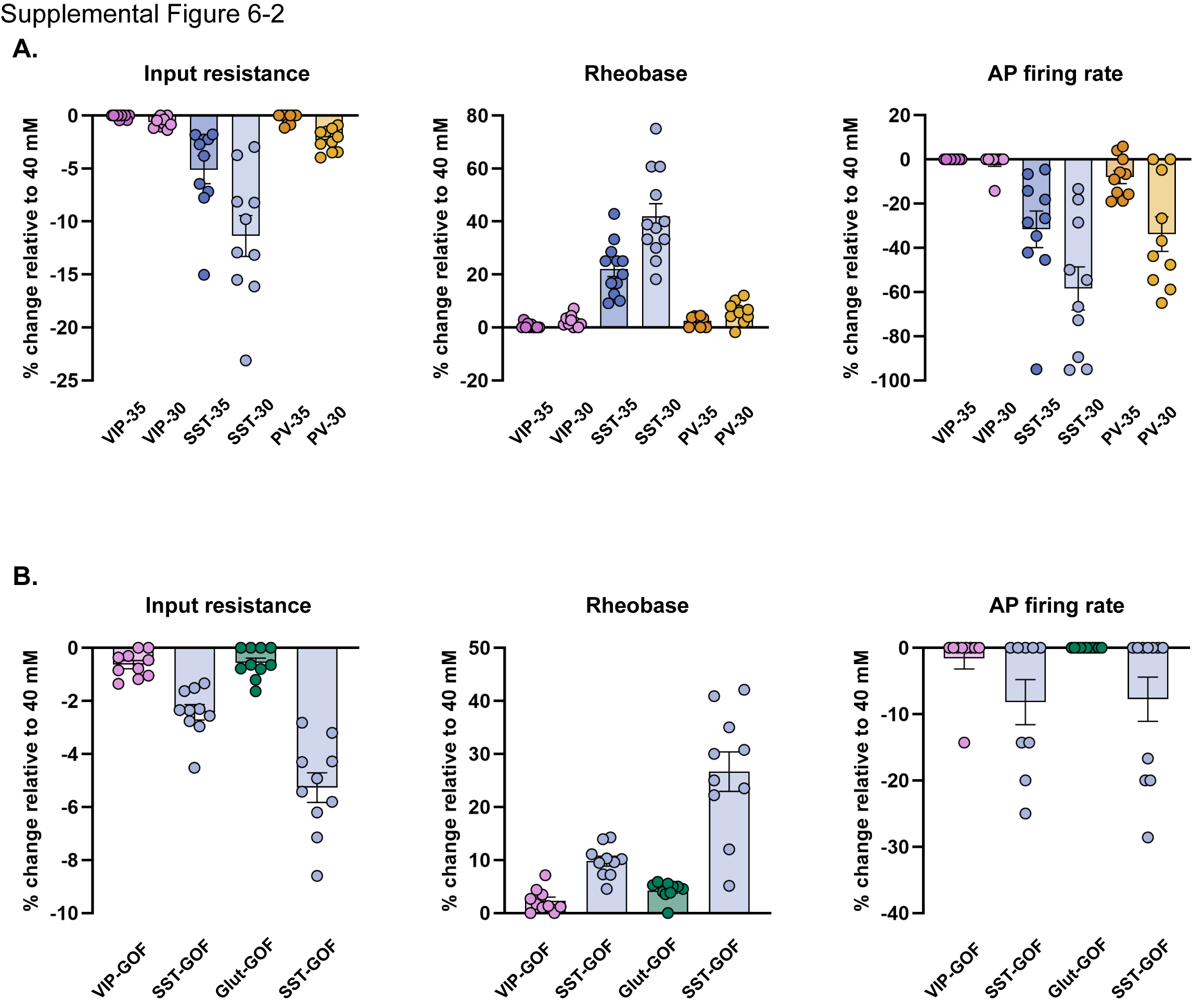

### Extended Figure 7-1

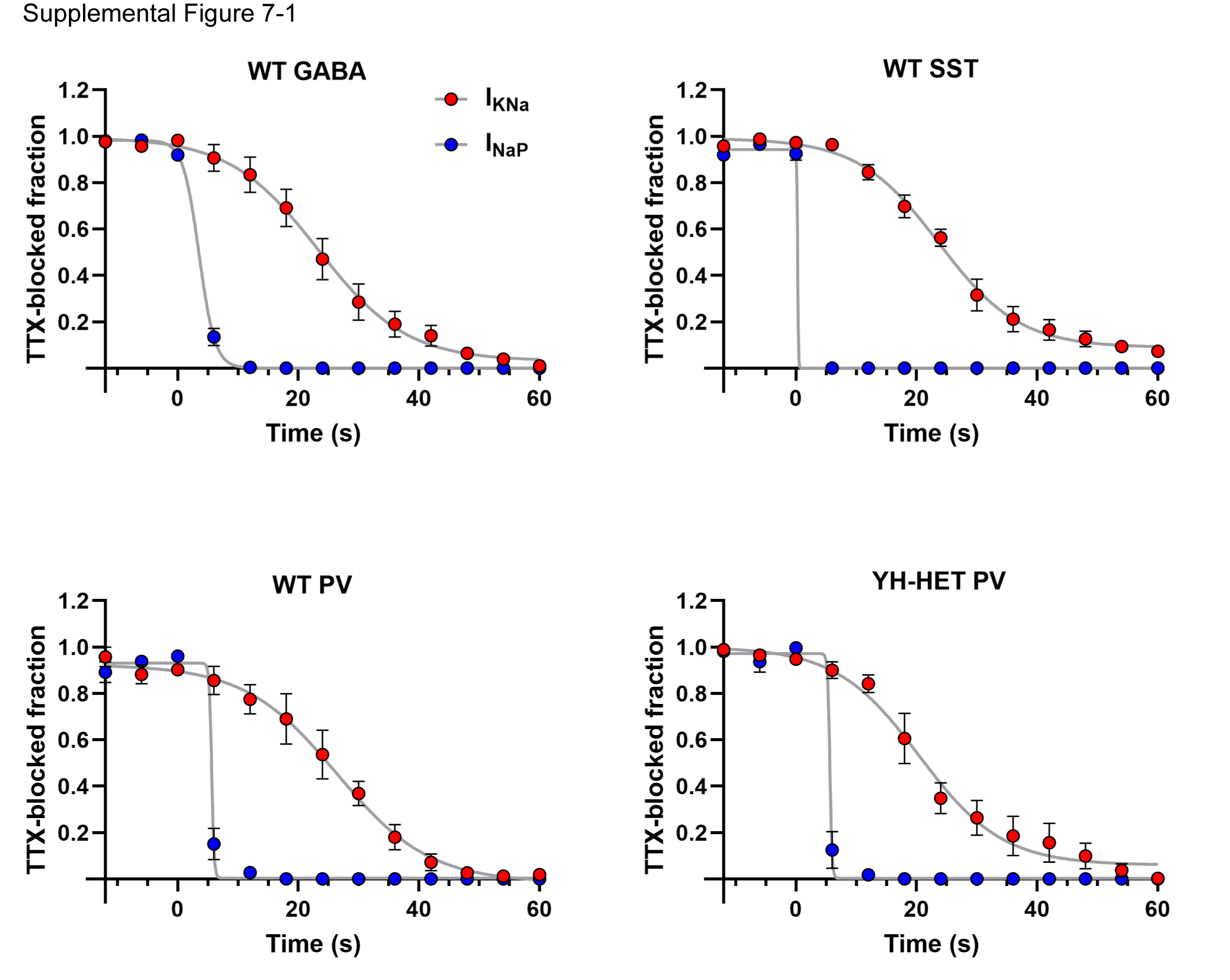

### Extended Figure 7-2

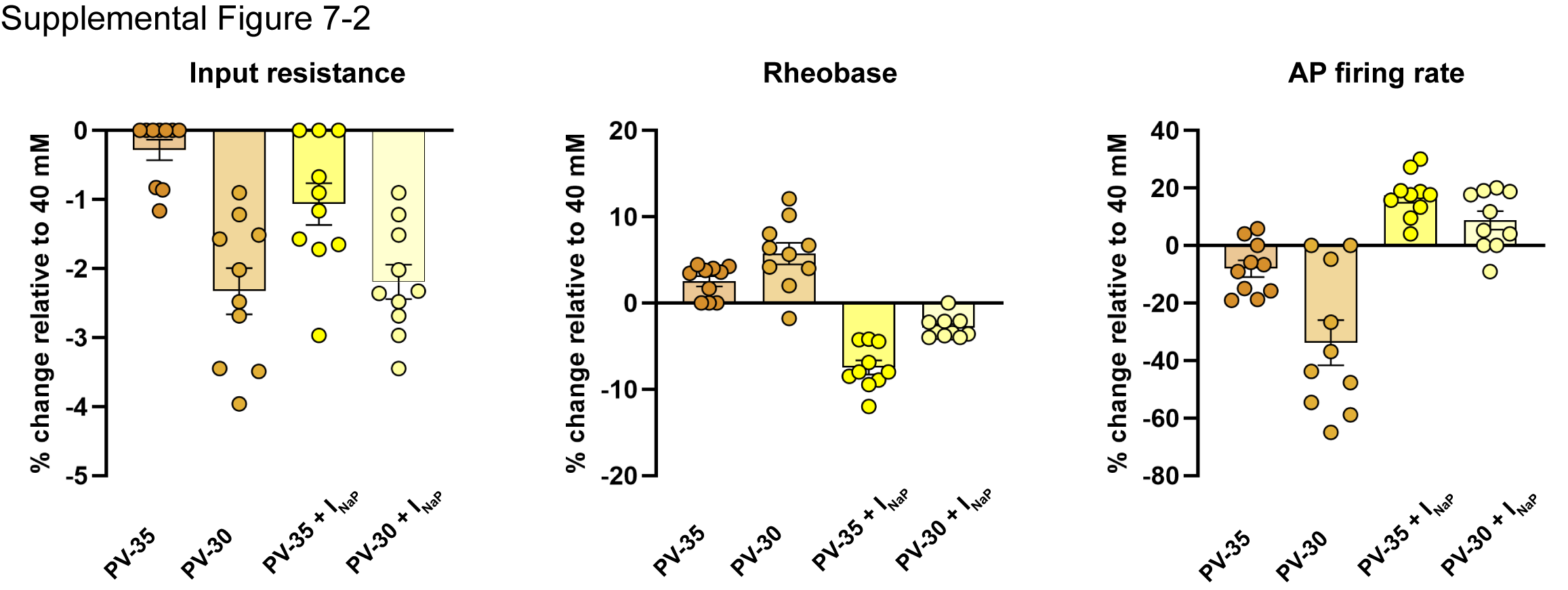
